# Testing the principles of Mendelian randomization: Opportunities and complications on a genomewide scale

**DOI:** 10.1101/124362

**Authors:** M Taylor, KE Tansey, DA Lawlor, J Bowden, DM Evans, Smith G Davey, NJ Timpson

**Author notes:** Corresponding Author: Nicholas Timpson. Current Address: School of Social and Community Medicine, University of Bristol, Oakfield House, Oakfield Grove, Bristol, BS8 2BN.

## Abstract

**Background:** Mendelian randomization (MR) uses genetic variants as instrumental variables to assess whether observational associations between exposures and disease reflect causal relationships. MR requires genetic variants to be independent of factors that confound observational associations.

**Methods:** Using data from the Avon Longitudinal Study of Parents and Children, associations within and between 121 phenotypes and 13,720 genetic variants (from the NHGRI-EBI GWAS catalog) were examined to assess the validity of MR assumptions.

**Results:** Amongst 7,260 pairwise comparisons between the 121 phenotypes, 2,188 (30%) provided evidence of association, where 363 were expected at the 5% level (observed:expected ratio=6.03; 95% CI: 5.42, 6.70; χ^2^=9682.29; d.f. =1, P≤1x10^-50^). Amongst 1,660,120 pairwise associations between phenotypes and genotypes, 86,748 (5.2%) gave evidence of association at the same threshold, where 83,006 were expected (observed:expected ratio=1.05; 95% CI: 1.04, 1.05; χ^2^=117.57; d.f. =1, P=2.15x10^-27^). Amongst 1,171,764 pairwise associations between the phenotypes and LD pruned independent genetic variants, 60,136 (5.1%) gave evidence of association, where 58,588 were expected (observed:expected ratio=1.03; 95% CI: 1.03, 1.08; χ^2^= 43.05; d.f. = 1, P=5.33x10^-11^).

**Conclusion:** These results confirm previously observed patterns of phenotypic correlation. They also provide evidence of a substantially lower level of association between genetic variants and phenotypes, with residual inflation the likely product of indistinguishable real genetic association, multiple variables measuring the same biological phenomena, or pleiotropy. These results reflect the favorable properties of genetic instruments for estimating causal relationships, but confirm the need for functional information or analytical methods to account for pleiotropic events.

---

Mendelian randomization (MR) is an analytical approach that uses genetic variants reliably associated with a risk factor of interest to estimate and quantify the causal relationship between exposure and a health related outcome (1, 2). The method exploits the properties of genetic variation and its validity rests upon three core assumptions: First, the genetic variation is associated with the exposure of interest; second, the genetic variation is independent of any factors that confound the association between the exposure and outcome; third, the genetic variation only influences the outcome via the causal effect of the exposure. In the form of instrumental variable (IV) analysis, these assumptions can be presented more formally; a genetic variant is a valid instrument for the purposes of estimating the causal effect of an exposure X on an outcome Y if it is: Associated with X (IV assumption 1); independent of all confounders and X and Y, denoted by U (IV assumption 2); and independent of Y given X and U (IV assumption 3) (**Figure 1**). Under these conditions, genetic variants associated with risk factors of interest are independent of other genetic variants and of phenotypes which might confound associations. Furthermore, as the outcome being measured cannot alter germline genotype, any associations observed between the genotype and outcome are unlikely to be affected by reverse causation and other traditional forms of study bias are likely to be avoided (1). If these assumptions are met then IV analyses (3) can be used to test for a causal relationship between exposure and outcome in observational studies and estimate the magnitude of this causal effect (4, 5).

**Figure 1.**
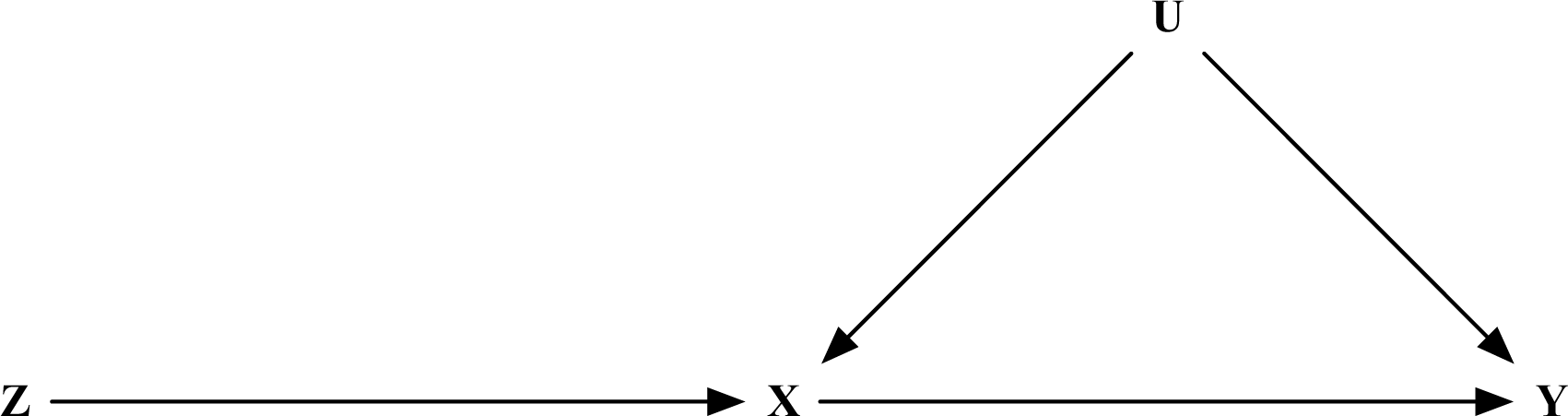
Directed acyclic graph representing the assumptions of instrumental variable analysis applied to Mendelian randomisation. Instrumental variable (IV) assumptions formally: a genetic variant is a valid instrument for the purposes of estimating the causal effect of an exposure X on an outcome Y if it is: Associated with X (IV assumption 1); independent of all confounders and X and Y, denoted by U (IV assumption 2); and independent of Y given X and U (IV assumption 3)

A previous study examined the validity of the assumption that there is no association between the genetic IV and confounders of the observational association by testing the association between 23 SNPs (selected on the basis of candidacy for phenotypic association) and 96 phenotypes in >4000 women from the British Women’s Heart and Health Study (6). Substantial pairwise association was found between phenotypes (i.e. 2036 (45%) associations from 4560 pairwise comparisons met a 1% level of statistical significance when only 45 would be expected by chance alone). In contrast, there was no substantial evidence for inflation in the number of associations between genetic and phenotypic variables (6). This study supported assertions as to the independence of genetic variation, but it did not use a large and comprehensive list of genetic variants known to be associated with complex traits/diseases that is now available (7) and increasingly used in MR studies.

Over the last decade, the rapid increase in the number of genome wide association studies (GWAS) has produced a large number of single nucleotide polymorphisms (SNPs) that are reliably associated with phenotypes and that might be used to test the causal effect of those phenotypes with disease outcomes. Here our aim was to use all SNPs listed in the NHGRI-EBI GWAS catalog to examine evidence against the assumptions underlying MR. To do this we systematically assessed pairwise associations between a broad collection of phenotypic characteristics that are often considered as outcomes, but also as confounding variables in observational studies and a large number of independent SNPs that are potential IVs in MR studies.

## METHODS

### Study participants

Genotypic and phenotypic data were from the Avon Longitudinal Study of Parents and Children (ALSPAC), a longitudinal study situated in the South West of England. ALSPAC recruited pregnant women between 1991 and 1992, with over 13,000 live births resulting from these pregnancies. Participants have been followed up with a series of questionnaires, clinic and lab-based assessments over the past 24 years, which has allowed for a wide range of phenotypic and biological measures to be collected. A total of 8,237 of the offspring were available for this study as only those participants with available genetic data (after quality control checks (8, 9)) were used. Ethical approval for the study was obtained from the ALSPAC Ethics and Law Committee and the Local Research Ethics Committees. Further information on the recruitment process is documented elsewhere (10, 11). The study website contains details of all the data that is available through a full searchable data dictionary: www.bris.ac.uk/alspac/researchers/data-access/data-dictionary/.

### Phenotypic measurements

The objective of this study was to assess a large set of phenotypes collected from a representative recruitment clinic of the ALSPAC study as a real-world example of a broad phenotypic data set. 121 phenotypic variables (a mixture of binary, continuous and categorical) were taken from clinic-based measurements and self-reported questionnaire data at approximately 7 years (10). This was the first face-to-face contact clinic undertaken within the ALSPAC study and has the largest participant response compared with all other clinic assessments to date. Large prospective cohorts, such as ALSPAC, often have hundreds of thousands of measured characteristics that are potential causal risk factors or confounders in epidemiological analyses. However, these represent a mixture of repeat measurements of the same characteristic (e.g. repeat weight, height and blood pressure at different ages) or measurements that reflect a similar characteristic (e.g. BMI, waist and directly assessed fat mass as measures of adiposity). Since we know that such repeat measurements and different measures of the same thing will be highly correlated we purposely focused on the one assessment at age 7. We did, however, enrich those data with data from prenatal questionnaires, such as family background socioeconomic position and parental behaviors. As in a previous investigation (6), phenotypes with a known relationship to another were not included, for example, we included diastolic blood pressure and therefore did not include systolic blood pressure. Previous sensitivity analysis demonstrated that the proportion of ‘statistically significant’ correlations did not change depending on which of these variables were used (6). The full list of phenotypes used in this study is provided in **Supplementary Table 1.**

### Genetic data and SNP selection

A set of single nucleotide polymorphisms (SNPs) identified by GWAS as reliably associated with complex phenotypes were obtained from the NHGRI-EBI GWAS catalogue (12) (formerly the NCBI GWAS catalog; accessed 10^th^ June 2015) and data form them retrieved from ALSPAC participants. These variants were chosen to provide an inclusive list of confirmed GWAS associations that are most likely to be used in MR experiments (13). We generated an initial list of 17,272 SNPs from 2,154 studies and after removal of duplicate SNPs and those not available in the ALSPAC cohort 14,911 SNPs remained. In ALSPAC data, SNPs were subsequently filtered to MAF>1% and imputation quality score (INFO) > 0.8. Following the removal of SNPs based on these criteria a total 13,720 SNPs remained. All SNPs were bi-allelic and coded 0, 1 and 2 and analyzed assuming an additive genetic model. Full information on genetic data and imputation in the ALSPAC cohort is provided in **Supplementary methods.**

We also created a dataset of SNPs that were pruned agnostically on the basis of linkage disequilibrium (LD) between them using the --indep-pairwise function in PLINKv1.9 (14). This function takes three parameters: (1) window size; (2) step size; and (3) r^2^ threshold. LD is calculated for all SNPs within a window (size set to 1500 base pairs) and one SNP in a pair is removed if r^2^>0.2 and the window moved along the chromosome by the “step size” (150 base pairs). This procedure is then repeated. We removed regions with known long range LD (15) generating an LD restricted dataset of 9,684 SNPs.

### Statistical analysis

P value summaries of associations within phenotype data and between SNP and phenotype data were derived using a relevant form of regression (linear, logistic or ordinal logistic). The expected number of associations with alpha values of 5%, 1% and 0.01% were obtained by calculating 5%, 1% and 0.01% of the number of pairwise associations tested, respectively. For all results, inverse log P values (observed and expected) were generated and qq plots demonstrating the relationship between the observed and expected p value distributions drawn. All statistical analysis was carried out using Stata 13 (16) (commands: *“regress”, “logistic”* and *“ologit”*).

### Sensitivity analysis

First, given their potential utility in MR studies, we restricted our pruned set of genetic variants to only those in the NHGRI-EBI GWAS catalog that meet genome-wide significance (P=5x10^-8^) and tested associations between these and our phenotypic variables as previously reported.

Second, to ensure that any associations observed between phenotypic and genotypic variants were not being inflated by unexpected genotype-genotype correlations, we calculated genotype-genotype correlations for ALSPAC after LD pruning. While biologically one would not expect genotype-genotype correlations of this nature, the presence of these through either mis-mapping, population stratification or chance could potentially complicate MR analyses. This is only a fundamental problem if the same associations were observed across multiple datasets and thus independent MR analyses. We therefore tested for these events in two additional datasets: Wellcome Trust Case Control Consortium (WTCCC) (17) and 1000 Genomes European sample (EUR) (18). This was done to determine if unexpected genotype-genotype correlations between LD-independent variants were consistent across different datasets or if they were unique. Information on genotyping, quality control and pruning for these two datasets can be found in **Supplementary methods.** Pearson’s correlations for all pairwise combinations of SNPs for each dataset were determined. This analysis was conducted using the R statistical package (19).

Finally, as a comparator phenotype data set with an enrichment for biomedical and serological measures within the same participants, analyses were undertaken using the same phenotypic variable set used in the UK10K consortium cohorts arm for the ALSPAC collection (20).

## RESULTS

### Study description

The sample size of the variables ranged from 1,895 to 6,494 and included a large variety of phenotypic measures including physiological, socioeconomic and behavioral phenotypes at age 7 (**Supplementary Table 1**). A total of 13,720 SNPs were included in the analysis. MAF of these SNPs ranged from 0.01 to 0.50. In the LD restricted dataset, the MAF of 9,684 SNPs ranged from 0.01 to 0.50.

### Associations between phenotypes

In a total of 121 phenotypes, 7,260 pairwise comparisons were possible. 2,188 (30%) gave evidence of association where 363 were expected given an alpha level of 5% (observed:expected ratio=6.03; 95% CI: 5.42, 6.70; χ^2^=9682.29; d.f. =1, P≤1x10^-50^). This excess of association was also seen at alpha levels of 1% and 0.01% (**Table 1** and **Figure 2**).

**Table 1.** Summary of phenotypic associations observed between all 121 phenotypes.

| Associations | Cut off $P$ value | Observed N (%) | Expected N (%) | $\chi^2$ (d.f, $P$ value) | O:E ratio ((95% CI), $P$ value) |
| --- | --- | --- | --- | --- | --- |
| 7260 | 0.05 | 2,188 (30.1) | 363 (5.0) | 9682.29 (1, $\leq 1 \times 10^{-50}$) | 6.03 ((5.42,6.70), $\leq 1 \times 10^{-50}$) |
| | 0.01 | 1,542 (21.2) | 73 (1.0) | 29861.37 (1, $\leq 1 \times 10^{-50}$) | 21.12 ((16.74,26.65), $\leq 1 \times 10^{-50}$) |
| | 0.0001 | 869 (12.0) | 7 (0.01) | 106251.60 (1, $\leq 1 \times 10^{-50}$) | 124.14 ((59.05,260.99), $\leq 1 \times 10^{-50}$) |

**Figure 2.**
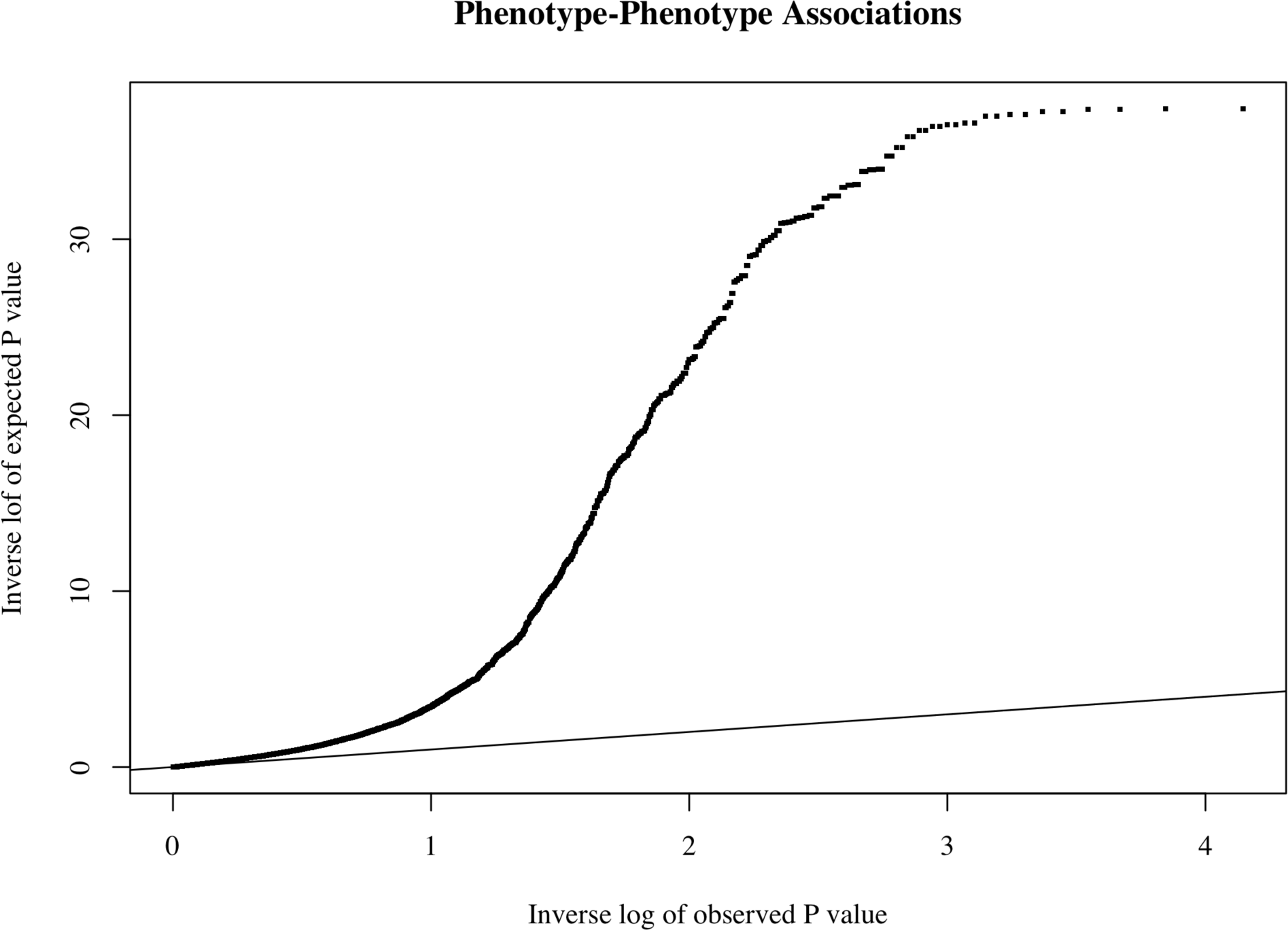
QQ plot of association between phenotypes. QQ plot of observed associations between 121 phenotypes included in analysis against expected P values. Inverse log of P values is the -log10 P value.

### Association between genetic variants and phenotypes

In a total of 13,720 SNPs and 121 phenotypes, 1,660,120 pairwise comparisons were possible. 86,748 (5.2%) gave evidence of association where 83,006 were expected given an alpha level of 5% (observed:expected ratio=1.05; 95% CI: 1.04, 1.05; χ^2^=117.57; d.f. =1, P2.15x10^-27^). Similar patterns of association were also seen at alpha levels of 1% and 0.01%. In a total of 9,684 SNPs (pruned for LD) and 121 phenotypes, 1,171,764 pairwise comparisons were possible. 60,136 (5.1%) gave evidence of association where 58,588 were expected (observed:expected ratio=1.03; 95% CI: 1.03, 1.08; χ^2^= 43.05; d.f. = 1, P=5.33x10^-11^) (**Table 2** and **Figure 3**).

**Table 2.** Summary Of Phenotypic and Genotypic Pairwise Associations Observed Between 121 Phenotypes And Genotypes (Full Set And Pruned)

| Dataset | Number of SNPs | Associations | Cut off $P$ value | Observed N (%) | Expected N (%) | $\chi^2$ (d.f, $P$ value) | O:E ratio ((95% CI), $P$ value) |
| --- | --- | --- | --- | --- | --- | --- | --- |
| All catalog SNPS | 13,720 | 1,660,120 | 0.05 | 86,748 (5.2) | 83,006 (5.0) | 117.57(1, $2.15 \times 10^{-27}$) | 1.05 ((1.04,1.05), $\leq 1 \times 10^{-50}$) |
| | | | 0.01 | 18,297 (1.1) | 16,601 (1.0) | 173.27 (1, $1.43 \times 10^{-39}$) | 1.10 ((1.08,1.13), $\leq 1 \times 10^{-50}$) |
| | | | 0.0001 | 366 (0.02) | 166 (0.01) | 240.99 (1, $\leq 1 \times 10^{-50}$) | 2.20 ((1.84,2.65), $\leq 1 \times 10^{-50}$) |
| LD pruned catalog SNPS | 9,684 | 1,171,764 | 0.05 | 60,136 (5.1) | 58,588 (5.0) | 43.05 (1, $5.33 \times 10^{-11}$) | 1.03 ((1.02,1.04), $4.01 \times 10^{-06}$) |
| | | | 0.01 | 12,388 (1.1) | 11717 (1.0) | 38.81 (1, $4.66 \times 10^{-10}$) | 1.06 ((1.03,1.08), 0.000014) |
| | | | 0.0001 | 184 (0.02) | 117 (0.01) | 38.37 (1, $5.85 \times 10^{-10}$) | 1.57 ((1.25,1.98), 0.00013) |

**Figure 3.**
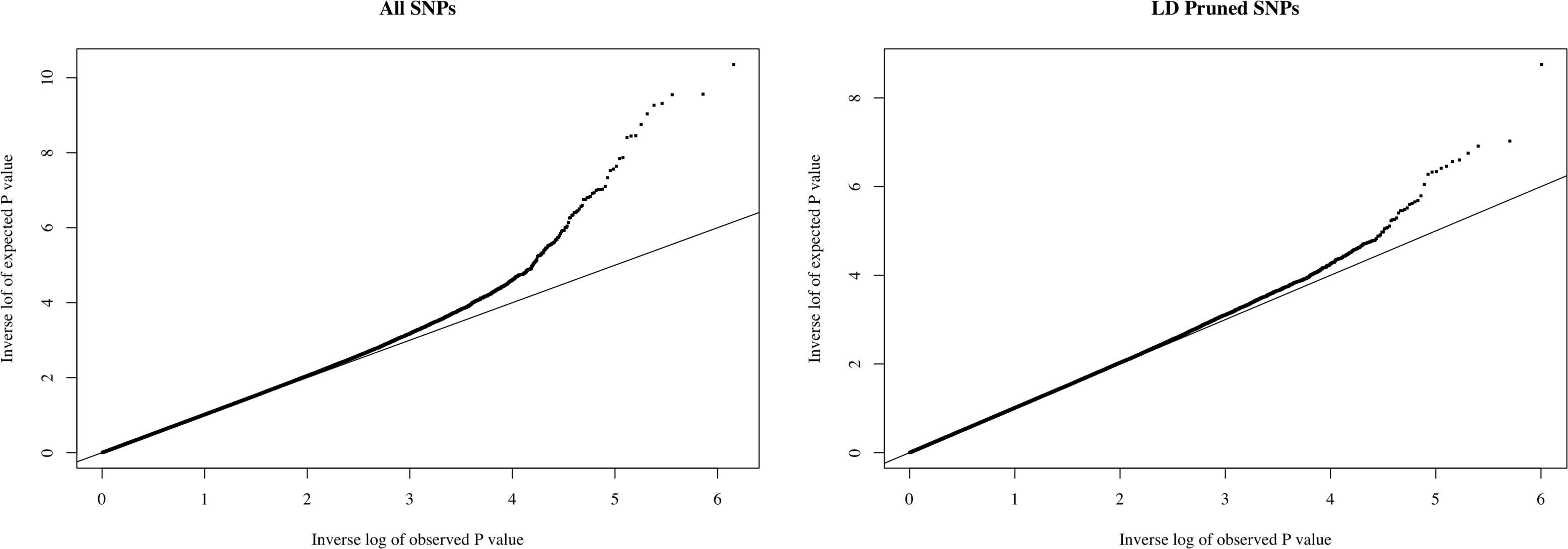
QQ plots of genotypic and phenotypic associations. Left: QQ plot of observed associations between 13,720 NHGRI-EBI-GWAS catalog SNPs (All SNPs) and 121 phenotypes included in analysis against expected P values. Right: QQ plot of observed associations between 9,684 NHGRI-EBI GWAS catalog SNPs pruned for LD and 121 phenotypes included in analysis against expected P values. Inverse log of P values is the -log10 P value.

As an indicator of genotype-phenotype associations which would have been followed up in bespoke GWAS efforts in the ALSPAC collection, associations with a P value below the GWAS threshold of P=5x10^-8^ were examined in further detail (**Supplementary Table 2**). rs9568856 (chromosome 13) was associated with both hip circumference and arm circumference. This genetic variant has previously been shown to be associated with body mass index (P = 7.75x10^-5^) and hip circumference (P = 1.7x10^-5^) in a large GWAS from The Genetic Investigation of Anthropometric Traits (GIANT) consortium (21, 22). Additionally, we observed an association between rs4820268 (chromosome 22) and with haemoglobin levels. This genetic variant has previous been shown to be associated with several haematological traits (23), including iron levels (P’s = 5.5x10^-7^ (24) and 2.4x10^-11^ (25)) and haemoglobin levels (P = 7.4x10^-7^ (26)). Of six additional associations, five were observed in strong LD with those identified above and the remaining observed association was between rs2290400 (chromosome 17, trait: Type I diabetes) and asthma. When using a Bonferroni corrected threshold calculated from the number of tests in this specific analysis (P=3.10^-8^), all but one (the association between rs2290400 and asthma) remained.

### Sensitivity analysis

When restricting our SNPs to those in the NHGRI-EBI GWAS catalog that were genome-wide significant, 6,175 SNPs were included in the analysis and results were consistent with those reported above (**Supplementary Table 3, Supplementary Figure 1**).

We quantified the proportion of genotype-genotype pairs with evidence for association within ALSPAC, EUR and WTCCC, then those pairs with evidence for association in two datasets and those that had evidence for association in all three datasets (**Supplementary Figure 2**). There was some evidence of inflated genotype-genotype correlations in ALSPAC, but only a small proportion of the total number of correlations within genetic variants retained evidence for association after correction for multiple testing (**Supplementary Table 4**) and very few were shared across all three datasets (0.003%).

Analysis undertaken using the same framework, but with an enrichment for biomedical and serological measures showed a relatively small inflation in the number of associations however results were consistent with those reported above (**Supplementary Table 5**). Results using the biomedical and serological measures only also followed the same patterns of association (**Supplementary Table 6**).

## DISCUSSION

By taking a wide selection of phenotypes along with GWAS catalog SNPs we have demonstrated the richness of phenotypic correlation and confirmed the comparatively sparse nature of broadly assessed genotype-phenotype association. More than 30% of pairwise associations between phenotypic characteristics in our dataset showed evidence for association, this empirically supporting the notion that observational epidemiological studies are potentially prone to confounding by physiological, socioeconomic and behavioral characteristics (27, 28). In contrast to this, only 5.1% of pairwise associations between phenotypic and genotypic characteristics gave evidence for association; a rate closely aligned to chance and likely inflated by real genotype-phenotype association or multiple variables measuring the same biological phenomena (poor measurement). This pattern of association between phenotypic traits and the contrasting pattern between genetic variants selected from GWAS and phenotypic traits was also observed at alpha values of 1% and 0.01% and reflects important properties of genetic variation relevant to causal analyses.

In a previous study, the number of observed associations was 11 times higher than the number of expected associations at the 5% alpha level (6). This is higher than the 6-fold difference observed in this study based on a more comprehensive data set. As the previous study was carried out on older women (aged between 60 and 79), this difference may reflect the fact that in general phenotypes become more correlated with increasing age as a result of exposure to overarching networks of environmental confounders. Despite this difference, our results highlight the marked inflation in correlation between phenotypic measurements and hence the difficulty in studies using multivariable regression approaches to be confident that residual or unmeasured confounding does not yield exaggerated or attenuated results (29).

This study provides support for the second assumption of MR analyses - that genotype is not associated with factors that confound the relationship between the exposure and the outcome – but has limitations. Despite patterns observed here, it remains important to check that genetic instruments are not related to observed potential confounders in MR studies; an event which could theoretically be driven by specific instances of population stratification or chance. In addition, this study was not explicitly designed to address the presence of horizontal/true pleiotropy, which is often analytically indistinguishable from vertical pleiotropy or pathway effects (30). It is possible that the observed inflation in the number of observed genotype/phenotype associations is predominantly driven by vertical/pathway based associations and results from the more biomedically aligned phenotype panel may be considered in line with this. However, we urge caution in MR analyses in the absence of testing this interpretation formally. Indeed, whilst the SNPs found in this study that might be considered as potentially valid instruments for MR analysis (i.e. those meeting genomewide significance, p=5x10^-8^) only had 1 trait association each, in more comprehensive analyses, shared genetic contributions to phenotypes of interest were prevalent (31). These extent of these types of event are of course subject to the statistical performance of the study in question, but are presented to illustrate that even with the contrasting scale of variable correlation in genotypic data, the underpinnings of applied genotypic data should be explored or accounted for.

We strongly support the use of methods attempting to account for events like genetic confounding (pleiotropy) that can distort causal estimates. This is especially true given the trend for using large collections of potentially invalid instruments in polygenic risk score based analyses (20). MR-Egger regression and the weighted median approach (32, 33) are promising developments and able to estimate causal effects in the presence of pleiotropic instruments (32). These approaches require additional assumptions about the nature of pleiotropy to be made and provide causal estimates with a reduced precision, however are likely to give refinement to this paradigm of natural experimentation (34) which might otherwise suffer from inevitable limitations of its own design. This serves as a reminder of the utility of MR as providing an important contribution to the evidence base as opposed to a definitive answer (35).

In conclusion, this study has reaffirmed our understanding of the extensive patterns of correlation seen between non-genetic traits. In a systematic manner and pertinent to the explosion of MR use and methods development currently, it has also shown that this is not the case for correlations between genetic variants and phenotypes. This provides further support for the use of genetic variants as IVs in MR studies as they are unlikely to generally strongly violate the assumptions that IVs are not related to confounders of the observational association.

## ACKNOWLEDGEMENTS

We are extremely grateful to all the families who took part in this study, the midwives for their help in recruiting them and the whole ALSPAC team, which includes interviewers, computer and laboratory technicians, clerical workers, research scientists, volunteers, managers, receptionists and nurses.

## REFERENCES

1. Davey Smith G, Ebrahim S. ‘Mendelian randomization’: can genetic epidemiology contribute to understanding environmental determinants of disease? International journal of epidemiology 2003;32(1):1–22.

2. Davey Smith G, Hemani G. Mendelian randomization: genetic anchors for causal inference in epidemiological studies. Human molecular genetics 2014;23(R1):R89–R98.

3. Lawlor DA, Harbord RM, Sterne JA, et al. Mendelian randomization: using genes as instruments for making causal inferences in epidemiology. Statistics in medicine 2008;27(8):1133–63.

4. Greenland S. An introduction to instrumental variables for epidemiologists. International journal of epidemiology 2000;29(4):722–9.

5. Angrist JD, Imbens GW, Rubin DB. Identification of causal effects using instrumental variables. Journal of the American Statistical Association 1996;91:444–72.

6. Davey Smith G, Lawlor DA, Harbord R, et al. Clustered Environments and Randomized Genes: A Fundamental Distinction between Conventional and Genetic Epidemiology. PLoS medicine 2007;4(12):e352.

7. Hindorff LA, MacArthur J, Wise A, et al. A Catalog of Published Genome-Wide Association Studies. (www.genome.gov/gwasstudies). (Accessed 02/05/2014).

8. Paternoster L, Howe LD, Tilling K, et al. Adult height variants affect birth length and growth rate in children. Human molecular genetics 2011;20(20):4069–75.

9. Sieradzka D, Power RA, Freeman D, et al. Are genetic risk factors for psychosis also associated with dimension-specific psychotic experiences in adolescence? PloS one 2014;9(4):e94398.

10. Boyd A, Golding J, Macleod J, et al. Cohort Profile: The ‘Children of the 90s’‐‐the index offspring of the Avon Longitudinal Study of Parents and Children. Int J Epidemiol 2012.

11. Fraser A, Macdonald-Wallis C, Tilling K, et al. Cohort Profile: the Avon Longitudinal Study of Parents and Children: ALSPAC mothers cohort. International journal of epidemiology 2013;42(1):97–110.

12. Welter D, MacArthur J, Morales J, et al. The NHGRI GWAS Catalog, a curated resource of SNP-trait associations. Nucleic acids research 2014;42(D1):D1001–D6.

13. Lawlor DA, Windmeijer F, Davey Smith G. Is Mendelian randomization ‘lost in translation?’: comments on ‘Mendelian randomization equals instrumental variable analysis with genetic instruments’ by Wehby et al. Statistics in medicine 2008;27(15):2750–5.

14. Chang C, Chow C, Tellier L, et al. Second-generation PLINK: rising to the challenge of larger and richers datasets. GigaScience 2015;4.

15. Price AL, Weale ME, Patterson N, et al. Long-range LD can confound genome scans in admixed populations. The American Journal of Human Genetics 2008;83(1):132–5.

16. StataCorp. Stata Statistical Software: Release 13. 2013.

17. Burton PR, Clayton DG, Cardon LR, et al. Genome-wide association study of 14,000 cases of seven common diseases and 3,000 shared controls. Nature 2007;447(7145):661–78.

18. The 1000 Genomes Consortium. A global reference for human genetic variation. Nature 2015;526(7571):68–74.

19. R Development Core Team. R: A language and environment for statistical computing. R Foundation for Statistical Computing, Vienna, Austria. 2013. ISBN 3-900051-07-0, 2014.

20. The UK10K Consortium. The UK10K project identifies rare variants in health and disease. Nature 2015;526(7571):82–90.

21. Speliotes EK, Willer CJ, Berndt SI, et al. Association analyses of 249,796 individuals reveal 18 new loci associated with body mass index. Nature genetics 2010;42(11):937–48.

22. Berndt SI, Gustafsson S, Mägi R, et al. Genome-wide meta-analysis identifies 11 new loci for anthropometric traits and provides insights into genetic architecture. Nature genetics 2013;45(5):501–12.

23. Vasquez L, Mann A, Chen L, et al. From GWAS to function: lessons from blood cells. ISBT Science Series 2015.

24. Pichler I, Minelli C, Sanna S, et al. Identification of a common variant in the TFR2 gene implicated in the physiological regulation of serum iron levels. Human molecular genetics 2010:ddq552.

25. Tanaka T, Roy CN, Yao W, et al. A genome-wide association analysis of serum iron concentrations. Blood 2010;115(1):94–6.

26. Ferreira MA, Hottenga J-J, Warrington NM, et al. Sequence variants in three loci influence monocyte counts and erythrocyte volume. The American Journal of Human Genetics 2009;85(5):745–9.

27. Davey Smith G, Ebrahim S. Epidemiology‐‐is it time to call it a day? International journal of epidemiology 2001;30(1): 1–11.

28. Davey Smith G, Ebrahim S, Lewis S, et al. Genetic epidemiology and public health: hope, hype, and future prospects. Lancet 2005;366(9495):1484–98.

29. Fewell Z, Davey Smith G, Sterne JA. The impact of residual and unmeasured confounding in epidemiologic studies: a simulation study. American journal of epidemiology 2007;166(6):646–55.

30. Paaby AB, Rockman MV. The many faces of pleiotropy. Trends in Genetics 2013;29(2):66–73.

31. Pickrell JK, Berisa T, Liu JZ, et al. Detection and interpretation of shared genetic influences on 42 human traits. Nature genetics 2016.

32. Bowden J, Davey Smith G, Burgess S. Mendelian randomization with invalid instruments: effect estimation and bias detection through Egger regression. International journal of epidemiology 2015;44(2):512–25.

33. Bowden J, Davey Smith G, Haycock PC, et al. Consistent estimation in Mendelian randomization with some invalid instruments using a weighted median estimator. Genetic epidemiology 2016;40(4):304–14.

34. Hingorani A, Humphries S. Nature’s randomised trials. The Lancet 2005;366(9501):1906–8.

35. Burgess S, Timpson NJ, Ebrahim S, et al. Mendelian randomization: where are we now and where are we going? International journal of epidemiology 2015;44(2):379–88.

36. Delaneau O, Marchini J, The 1000 Genomes Consortium. Integrating sequence and array data to create an improved 1000 Genomes Project haplotype reference panel. Nature communications 2014;5.

37. Howie BN, Donnelly P, Marchini J. A flexible and accurate genotype imputation method for the next generation of genome-wide association studies. PLoS genetics 2009;5(6):e1000529.

38. Howie B, Marchini J, Stephens M. Genotype imputation with thousands of genomes. G3: Genes, Genomes, Genetics 2011;1(6):457–70.

39. Purcell S, Neale B, Todd-Brown K, et al. PLINK: a tool set for whole-genome association and population-based linkage analyses. The American Journal of Human Genetics 2007;81(3):559–75.

40. The 1000 Genomes Consortium. An integrated map of genetic variation from 1,092 human genomes. Nature 2012;491(7422):56–65.

